# Signatures of insecticide selection in the genome of *Drosophila melanogaster*

**DOI:** 10.1101/287250

**Authors:** David Duneau, Haina Sun, Jonathan Revah, Keri San Miguel, Henry D. Kunerth, Ian V. Caldas, Philipp W. Messer, Jeffrey G. Scott, Nicolas Buchon

## Abstract

Resistance to insecticides has evolved in multiple insect species, leading to increased application rates and even control failures. Understanding the genetic basis of insecticide resistance is fundamental for mitigating its impact on crop production and disease control. We performed a GWAS approach with the *Drosophila* Genetic Reference Panel (DGRP) to identify the mutations involved in resistance to two widely used classes of insecticides: organophosphates (OPs, parathion) and pyrethroids (deltamethrin). Most variation in parathion resistance was associated with mutations in the target gene *Ace*, while most variation in deltamethrin resistance was associated with mutations in *Cyp6a23*, a gene encoding a detoxification enzyme never previously associated with resistance. A “nested GWAS” further revealed the contribution of other loci: *Dscam1* and *trpl* were implicated in resistance to parathion, but only in lines lacking *Wolbachia*. *Cyp6a17*, the paralogous gene of *Cyp6a23*, and *CG7627*, an ATP-binding cassette transporter, were implicated in deltamethrin resistance. We observed signatures of recent selective sweeps at all of these resistance loci and confirmed that the soft sweep at *Ace* is indeed driven by the identified resistance mutations. Analysis of allele frequencies in additional population samples revealed that most resistance mutations are segregating across the globe, but that frequencies can vary substantially among populations. Altogether, our data reveal that the widely used OP and pyrethroid insecticides imposed a strong selection pressure on natural insect populations. However, it remains unclear why, in *Drosophila*, resistance evolved due to changes in the target site for OPs, but due to a detoxification enzyme for pyrethroids.

**Article summary:** Insecticides are widely used to control pests and insect vectors of disease. In response to the strong selection pressure exerted by insecticides, resistance has evolved in most insect species. We identified few genes present in several *Drosophila melanogaster* natural populations implicated in the evolution of resistance against two insecticides widely used today. We identified primary and secondary genes involved in the resistance. Surprisingly, resistance evolved in the target site for one insecticide, but was associated to changes in a novel detoxification enzyme for the other insecticide.

## Introduction

Insecticides are widely used for control of agricultural and structural pests, and to control insect vectors of disease. It is difficult, or perhaps impossible, to exactly calculate the economic and human health benefits associated with insecticide use, but they are significant. For example, depending on the crop and level of insect pressure present in a given year, insecticides can boost yields by 6-79% (Ware and Whitacre 2004). In just the USA, insecticide expenditures are >$6 billion and >550 million pounds are used annually (Meister and Sine 2014). In response to the strong selection pressure exerted by insecticides, resistance has evolved in multiple species against numerous insecticides. This can lead to increasing frequency of insecticide applications, increased application rates and even control failures; impacting both crop production and control of human (and animal) diseases. Thus, understanding the genetic basis underpinning the evolution of resistance to insecticides is of fundamental importance.

For more than twenty years, the availability of molecular tools has facilitated the identification of mutations responsible for changes in protein structure and also in gene expression causing insecticide resistance. Out of necessity these studies were usually carried out on strains that had been selected in the laboratory, in an effort to make the resistance gene(s) homozygous. Identification of the mutations responsible for resistance allowed for the frequency of these mutations to be examined in field populations. In the postgenomic era, Genome-Wide Association Studies (GWAS) offer the potential to examine how the evolution of insecticide resistance occurs at a whole genome level, without having to select a resistant strain in the laboratory. GWAS studies have been recently used to look at the pattern of resistance to a banned insecticide, (DDT, which has not been used in the USA since 1972), an organophosphate (OP, azinphos-methyl)) and a neonicotinoid insecticide (imidacloprid) (Battlay *et al*. 2016; Schmidt *et al*. 2017; Denecke *et al*. 2017), but have not yet been used to evaluate resistance to insecticides that have been and continue to be widely used, such as pyrethroids.

OP and pyrethroid insecticides are widely used today. OPs were developed in the late 1940s and were the most widely used class of insecticides for more than three decades. Pyrethroid insecticides were commercialized in the 1980s and rapidly replaced OPs as the most widely used class of insecticides for about 20 years. A great deal has been learned about the basis of resistance to these two classes of insecticides. Mutations in the target site (*acetylcholinesterase* also known as *Ace* or *AChE* for OPs and *voltage sensitive sodium channel* or *Vssc* for pyrethroids) and increased detoxification by cytochrome P450s [CYPs] and esterases/hydrolases are the major mechanisms of resistance (Newcomb *et al*. 1997; Scott 1999, 2017; Gunning and Moores 2001; Kono and Tomita 2006; Achaleke *et al*. 2009; Dong *et al*. 2014). Resistance due to increased detoxification is most commonly due to increased expression of a gene, but non-synonymous mutations can cause resistance as well. Understanding the role of metabolism in insecticide poisoning has been less clearly resolved than target site mutations because there are multiple potential detoxification protein families (CYPs, GSTs, esterases/hydrolases, etc.) and each of these groups of proteins contains multiple genes (e.g. often >100 *Cyp*s).

The aim of this study was to investigate the variation in resistance of individuals collected from a field population towards two classes of currently used insecticides in a natural population of *Drosophila melanogaster* using an unbiased approach able to reveal resistance loci (and candidate genes) in the whole genome. To this purpose, we performed GWAS using the *Drosophila* Genetic Reference Panel (DGRP), a panel of 205 lines of *D. melanogaster* mostly homozygous and fully sequenced and derived from a wild caught population (Mackay *et al*. 2012). The use of inbred fly lines allowed us to assess the impact of pesticides on distinct, but constant genetic backgrounds to tease out the effect of the genotype from environmental effects. The association of a particular allele at a particular locus with the degree of resistance of each line to an insecticide allowed us to identify candidate genes belonging to the quantitative trait loci (QTL) underlying the resistance to those insecticides. Using an approach that first performed a GWAS with all the *Drosophila* lines of the panel followed by another GWAS including only the lines that did not carry the major effect allele (nested GWAS), we were able to identify and validate a set of genes of major and minor effect on resistance to OPs (parathion) and to pyrethroids (deltamethrin). These classes of insecticides were selected because they have been widely used for decades and are representatives of the 3^rd^ and 2^nd^ most widely used classes of insecticides today (OPs and pyrethroids, respectively). We thus expected these pesticides to have exerted significant selection pressure on *D. melanogaster*. Using other *Drosophila* genetic panels, we investigated the presence of our detected mutations in other natural populations and evaluated the signal of selection on our detected mutations.

## Materials and Methods

### Fly stock and husbandry

All *Drosophila* stocks were raised at 22°C on standard Cornmeal agar medium, with a relative humidity of 60%-70%, and a photoperiod of 12L:12D, unless specified. For the Genome Wide Association Study (GWAS), most of the isogenic lines of the *Drosophila* Genetic Reference Panel were used (193 lines were exposed to parathion and 191 to deltamethrin) (Mackay *et al*. 2012; Huang *et al*. 2014). To evaluate the involvement of candidate genes in resistance, *UAS*-controlled *in vivo* RNAi and overexpression experiments were performed using either the *Actin5c-Gal4* driver (*Act5c-Gal4*) or the *da-Gal4; ubi-Gal80^TS^* conditional driver (*da-Gal4^TS^*). F_1_ progeny was obtained by crossing virgin females (25 isolated within 8 h of emergence) of the driver strain with males (∼15) of the *UAS-transgene* line. The F_1_ progenies (for crosses with the *Gal4^TS^ driver > UAS-transgene*) were raised at 18°C until three days after emergence, and then switched to 29°C for a week to trigger maximum transgene expression before being assayed for resistance to deltamethrin at 29°C. The F_1_ progenies (for crosses with the *Act5c-Gal4 driver > UAS-transgene*) were raised and assayed for resistance at 25°C. As a control, the driver virgin females were crossed to the appropriate background lines *Attp2*, *Attp40* or *w^1118^* (see Table S1).

**Table 1:** Genetic variation and heritability of susceptibility to two insecticides

| **Insecticides** | ***N* flies** | ***N* lines** | **$V_e$** | **$V_g$** | **$h^2$** |
| --- | --- | --- | --- | --- | --- |
| **parathion** | 194 | 16.568 | 6.04 | 43.83 | 0.88 |
| **deltamethrin** | 195 | 16.684 | 4.4 | 7.07 | 0.61 |

Thirty-three transgenic *Drosophila* lines and the appropriate background lines were obtained from the Bloomington *Drosophila* Stock Center (BDSC, Indiana University, Bloomington, IN, USA) and the Vienna *Drosophila* Ressource Center (VDRC) (Table S1). Three mutant lines, four transgenic *UAS-RNAi* lines and one overexpression line from the parathion candidate gene list were available for knockout, knockdown or overexpress of *trpl*, *olf413*, *fru* or *Dscam1* genes. One mutant line and nine transgenic *UAS-RNAi* lines from the deltamethrin candidate gene list were used for knockout of *Cyp6a17* or knockdown of *Cyp6a9*, *Cyp6a17*, *Cyp6a19*, *Cyp6a20*, *Cyp6a22*, *Cyp6a23*, *CG7627* and *tou*, respectively.

### Insecticides and bioassays

The residual contact application method was used to examine the relative susceptibility of DGRP lines for the insecticides, parathion and deltamethrin. Parathion (99.3%, Chem Service, West Chester, PA, USA) and deltamethrin (100%, Roussel UCLAF, Paris, France) were each dissolved in acetone to final concentrations of 1.5 µg/ml and 0.7 µg/ml respectively. 0.5 ml insecticide solution was added to a 38.6 cm^2^ scintillation vial (Wheaton Scientific, Millville, NJ, USA), which was coated evenly on the inside surface using a hotdog roller machine (Gold Medal, Cincinnati, OH, USA) for 20 min under a fume hood until all the acetone had evaporated. Treated vials were incubated at 23℃ for 20 hours before flies were transferred inside. Approximately 20 5-8 days old adult males for each line were assayed per vial for each insecticide. Vials were stoppered with a piece of cotton covered with a square of nylon tulle fabric and secured with a staple. The stopper was injected with 2 ml of 20% sugar water after addition of the flies, and assays were held at 25℃ with a photoperiod 12L:12D. 1 ml of distilled H_2_O was added to the stoppers after 24 h. For GWAS, mortality was assessed at 2.5 h, 5 h, 11 h, 24 h, and 48 h after flies were added to each vial for parathion and at 48 h for deltamethrin. Ataxic flies were counted as dead and five separate experiments were conducted over five continuous weeks. For validation experiments, mortality was assessed 24 h after insecticide treatment. F_1_ males (3-7-day-old) from each of the crosses were tested using single dose assays for parathion or deltamethrin.

### Genome wide association analysis

The genetic diversity of the DGRP lines comprises about 4 millions SNPs. However, the genotypic information for each line differs between loci (e.g. some loci have information for all lines, other do not), thus, sample sizes used in each association tested changes from a locus to another. Not all SNPs are therefore suitable for testing the association between the genetic variation at one locus and the resistance to insecticide. We selected SNPs for our association study based on 2 criteria: 1-avoid a complete collinearity (possibly confounding) between alleles and *Wolbachia* status (i.e. we excluded cases where one allele corresponds to *Wolbachia* infection and the other to an uninfected status); 2- we had enough lines per treatment to run the model. Prior to each test, we therefore calculated a two-by-two matrix with *Wolbachia* status and allele identity (i.e. W^+^/allele1, W^-^/allele1, W^+^/allele2, W^-^/allele2) summarizing the sum of lines for each category. We further included in our association only the SNPs where at least three of the categories had five lines. All the analyses were performed with custom made script.

We next estimated the significance of the alleles at each selected SNP for the survival of each line to parathion and deltamethrin. For parathion, we used a parametric survival analysis with a log-normal distribution of the error (Function Survreg from the R package “Survival”). The model was as following: Surv (Hour_of_death, Censor) ∼ *Wolbachia* status * SNP + frailty (Experiment, distribution=’gaussian’) + frailty (DGRP_lines, distribution=’gaussian’). The variable “Experiment” and the identity of the lines were accounted for as random effect following a Gaussian distribution. For the second insecticide, deltamethrin, we tested with a linear regression based on a binomial distribution of the error (function GLMER from the R package “lme4”), the survival at 48h post-exposure of the individuals carrying each allele. We could not use a survival analysis because between 2.5h and 48h some ataxic individuals could recover (temporally) before eventually dying. Therefore, the model was as following: cbind (Delta_alive, Delta_dead) ∼ *Wolbachia* + SNP +(1|DGRP_lines). The identity of the lines was accounted for as a random effect following a Gaussian distribution. We compared this analysis to the analysis accounting for the variable “Experiment” as a random effect. The results were not strongly different but the approach including a random effect required much more computer time (month of analysis instead of days). Therefore, we performed our analyses without this term. To identify other genes responsible for the resistance in absence of major effect alleles, we performed a “Nested-GWAS” which consists in running the same analysis on the lines that are not 100% survival. In other words, we attempted to find the alleles responsible for the remaining variation.

Candidate SNPs were among the alleles where the p-value was below 0.0001. We then converted the positions provided for the version 5 of the *D. melanogaster* genome annotation in version 6 with the convert tool from Flybase. The effect and the characterization of the mutation’s effect at each candidate SNP were provided using VEP from the website Ensembl (http://www.ensembl.org/info/docs/tools/vep/index.html). Candidates to be validated were chosen based on the shape of the peak in the Manhattan plot and the function provided by VEP (likelihood to be involved in the resistance). Then, those with a non-synonymous mutation were favored.

Validation of selected candidates were tested by exposing the genotypes and their control to the same conditions as in the GWAS. Differences of proportion of surviving individuals 48 hours post exposure were statistically tested with a generalized linear model with a quasibinomial distribution of the error. We used a general linear hypothesis test (glht) with Tukey post Hoc pairwise comparisons (alpha=0.05), to ascertain differences between pairs of treatments (package *multcomp* in R).

### Correlation of resistance with gene expression and other phenotypes known in the DGRP lines

To determine whether the resistance to each of the insecticide correlated with resistance to other abiotic stress such as paraquat, starvation and ethanol, we used measurement from other studies (Mackay *et al*. 2012; Weber *et al*. 2012; Morozova *et al*. 2015) and assessed the correlation (of Spearman) with our proportion of survival to our insecticides 48h post-exposure. We also tested whether the constitutive expression of our genes involved in resistance correlated with the resistance to pesticide. Although this approach is very limited as both phenotypes were obtained in different laboratories, we used the constitutive gene expression of our genes from (Huang *et al*. 2015) to correlate (Spearman) it with the proportion of survival individuals 48 hours post-exposure to the insecticides.

### Population genetic analyses

For the H12 selection scans and haplotype trees presented in Figure 4 we used VCF files from the DGRP 2 Freeze 2.0 calls (http://dgrp2.gnets.ncsu.edu/data.html). Only the lines that were included in the GWAS analysis were used. We further filtered out any site with more than 18% missing data. Indels were removed and the data was subset to biallelic sites. Missing data was imputed and remaining heterozygous sites were phased with Beagle 4.1, using windows of 50,000 sites and 15 iterations per window (Browning and Browning 2016). Each autosomal arm was scanned using the H12 script obtained from the SelectionHapStats repository provided in (Garud *et al*. 2015), using window sizes of 800 segregating sites. We extracted 200 kilobase genomic windows centered on the *Ace* and *Cyp6a23* gene positions from the DGRP data, as well as from two random genomic regions not associated with GWAS hits. These windows contained between 6000 and 8500 biallelic SNPs. For each window, we first calculated a distance matrix using the observed number of nucleotide differences in our filtered data set. From these distanced matrices we estimated neighbor-joining trees (Saitou and Nei 1987). At the *Ace* and *Cyp6a23* windows, individuals were classified according to presence (“1”) or absence (“0”) of individual insecticide resistance mutations (3R:13,243,332, 3R:13,243,686 and 3R:13,243,999 at Ace; 2R:14,876,125 and 2R:14,876,857 at Cyp6a23). Trees were estimated and drawn using the R package ape (Paradis *et al*. 2004). The specific midpoints of the four windows used for the trees in Figure 4A and the number of SNPs in each window are: (i) 2L:17,403,824, 7722 SNPs; (ii) 2R:14,876,125, 7726 SNPs; (iii) 3L:14,419,400, 8531 SNPs; (iv) 3R:19,817,445, 6141 SNPs.

Allele frequency estimates reported in Figure 4C were obtained from the same DGRP data set used for the H12 scans and haplotype trees, except that here we included indels because the resistant allele at CG7627 is a deletion. For the GDL lines, VCF files were obtained from the Clark Lab at Cornell University. Indel information was obtained from VCF files downloaded from the Poole Lab website (http://www.johnpool.net/genomes.html). The same 18% missing data filter was applied prior to imputation, and the remaining sites were again phased using Beagle 4.1, using windows of 50,000 sites and 15 iterations per window (Browning and Browning 2016).

### Data availability

*Drosophila* lines are listed in table S1 with their stock number. Raw phenotypic data and results from the GWAS are available in Supplemental Table S2, S3, S4, S5, S8 and S9.

## Results

Our results indicate that the resistance to an OP and pyrethroid in the DGRP lines is largely due to a single major locus, that additional loci provide minor effects, and that these loci differ between parathion and deltamethrin. Most variation in parathion resistance is associated with mutations in *Ace,* the target site of OPs (and carbamates). Most variation in deltamethrin resistance is associated with *Cyp6a23,* a probable detoxification enzyme. Both major effect genes were found under selection and we identified traces of soft sweep around their loci. Importantly, the alleles of the major effect genes we identified were not a particularity of our sampled population but were found in two other wild-caught *D. melanogaster* populations present in the Global Diversity panel lines (Grenier *et al*. 2015). Our study, therefore, reveals the specific and conserved mechanisms of resistance to various insecticides. Nested GWAS with the lines that did not carry the alleles 12 responsible for the major effects allowed us to identify the lesser contribution of other genes in the genome. We identified and validated the involvement of *Down syndrome cell adhesion molecule 1* (*Dscam1*) and *transient receptor potential-like* (*trpl*) in the resistance to parathion, and of *Cyp6a17* and *CG7627*, an ATP-binding cassette transporter in the resistance to deltamethrin.

### Genetic variation in insecticide resistance

To identify genes underlying natural variation in resistance to OPs and pyrethroids, we quantified the survival of DGRP lines to parathion and deltamethrin (194 lines for parathion and 195 for deltamethrin). Survival to parathion was monitored at 2.5 h, 5 h, 11 h, 24 h and 48 h post-exposure and the susceptibility of each line was estimated by comparing the time death took to happen among lines. For deltamethrin we could not monitor the time death took to happen because flies were ataxic early in the process but could sometimes recover before dying. Thus, we only monitored the proportion of dead individuals 48 h post-exposure (*i.e.* when ataxia was not a confounding effect anymore). The proportions of survival 48 h post-exposure were compared between lines for deltamethrin. We found striking and reproducible variation in the DGRP lines’ survival to both insecticides (Figure 1A).

**Figure 1.**
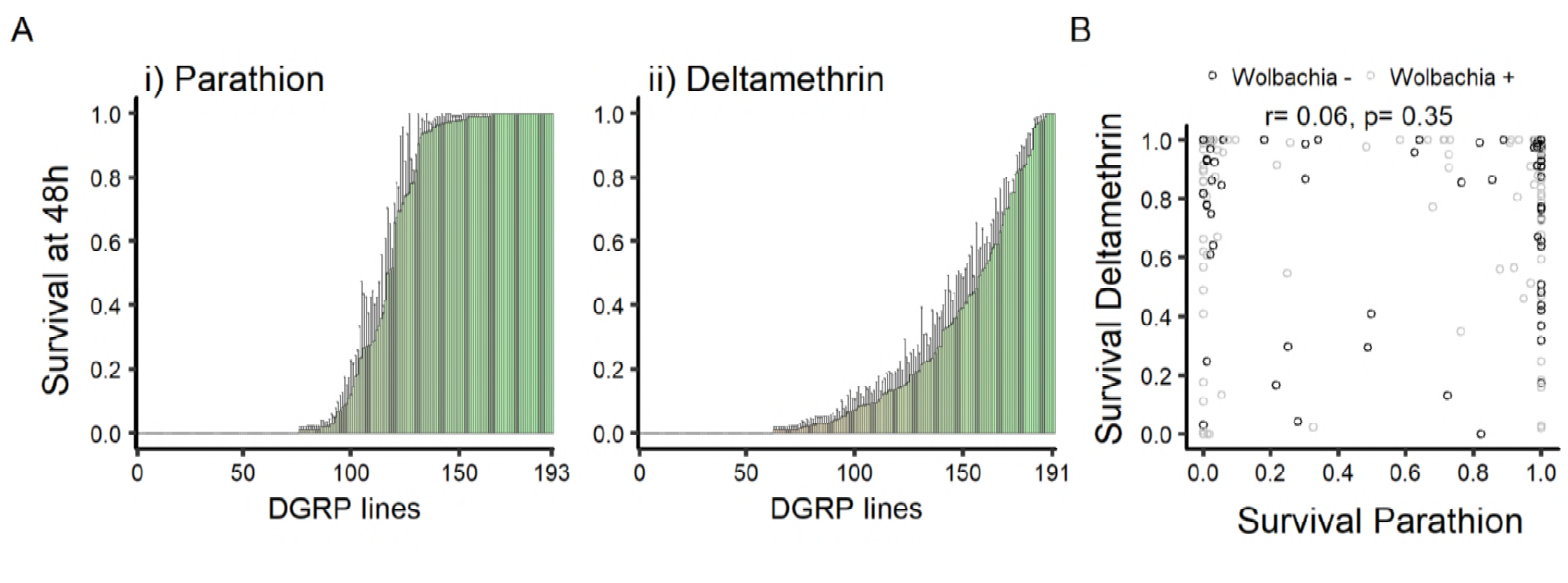
**A**-Ranked mean (± standard error) of male proportion surviving 48 h post-exposure to *i*) parathion and *ii*) deltamethrin. **B**-Correlation between resistance to parathion and resistance to deltamethrin. The resistance to one insecticide was not correlated to the resistance to the other insecticide. Analysis of correlation was done with Spearman correlation test.

Before examining the loci linked to resistance we investigated the role of non-genetic causes of differences in survival between the DGRP lines. Approximately half of the DGRP lines carry the bacterial endosymbiont *Wolbachia*. Therefore, we evaluated the possible contribution of *Wolbachia* to insecticide susceptibility with the average survivorship at each time point (Figure S1). Infection with *Wolbachia* did not correlate with resistance to parathion (Figure S1A) nor to resistance to deltamethrin (Figure S1B). Because resistance to different abiotic stresses could have shared mechanisms, we tested the correlations between resistance to parathion or deltamethrin and these stresses; namely the resistance to paraquat, starvation and ethanol that were measured in other studies (see details in methods, Figure S2). We did not detect any correlations with resistance to parathion. However, resistance to deltamethrin in our study correlated positively with both resistance to paraquat (r=0.18, p-value= 0.02) and resistance to starvation (r=0.25, p-value= 0.0004). Further studies would be needed to investigate these correlations, particularly because they were performed in different laboratories at different times. We next asked whether the variation we observed was due to genetic or environmental differences. The variation in insecticide resistance in our population was explained more by genetic variance than by environmental variance, with 88% heritability for sensitivity to parathion and 61% for deltamethrin (see Table 1). As DGRP lines show a high degree of genetic relatedness, it is possible that resistance to insecticides is an indirect consequence of physiological differences between lines. Thus, we next evaluated whether susceptibility to insecticide could be a secondary consequence to general physiological weakness of susceptible lines. To determine this, we compared the relative survival of individual DGRP lines to deltamethrin and parathion. The resistance to one insecticide was not correlated to the resistance to the other insecticide, suggesting that the determinants of resistance are not due to a simple resistance to stress and are specific to each insecticide (Figure 1B). In addition, individuals susceptible to insecticides were not more closely related among each other for either of the compounds tested (Figure S3).

Having ruled out non-genetic influences on survival to the insecticides, we next sought to identify the genetic determinants underlying variation in resistance to either parathion or deltamethrin. The ranked survival for parathion suggested a major allele effect due to the steep change in survival between lines (few lines are intermediates, Figure 1A*i*). However, the smooth continuum in the ranking of survival to deltamethrin (*i.e.* from lines that had 0% to 100% survivorship) suggested multiple loci could be involved in resistance (Figure 1A*ii*). We next estimated which loci could contribute to insecticide resistance by statistically associating mortality with the allelic polymorphism at each sequenced locus in the genome.

### Genetic basis of the variation in resistance to parathion

We first identified loci associated with resistance to parathion using GWAS. We tested the association of resistance to parathion with 1,784,231 SNPs/indels. In total, 44 loci were significantly associated (*i.e.* -log_10_(p-value) > 8) with resistance to parathion (Figure 2), but other SNPs/indels, less strongly associated, could be considered as candidates (271 had -log_10_(p-value) > 5 and 787 had -log_10_(p-value) > 4). The presumptive genetic alterations and consequences for the genes close to these SNPs/indels can be found in Table S2. Based on both the significance of the association (*i.e.* the peaks in the Manhattan plots, Figure 2) and the consequence of the genetic change associated with the SNPs/indels (priority to SNPs/indels altering protein structure or in introns/promoters based on prediction on the Ensembl website), we made a list of loci and built a list of genes likely to be involved in parathion resistance (black p-values in Figure S4). The most significant QTLs were located in *Ace* (Figure 2A). These QTLs were mapped to SNPs that generate non-synonymous mutations [F368Y in position 3R:13,243,332: Figure 2B*i*); G303A in position 3R:13,243,686: Figure S5A; I199V in position 3R:13,243,999: Figure S5B] in *Ace.* Previous work has shown these mutations confer resistance to organophosphates (Fournier *et al*. 1993). We therefore conclude that in the case of parathion resistance, variation in the target protein is responsible for most of the variation in resistance.

**Figure 2.**
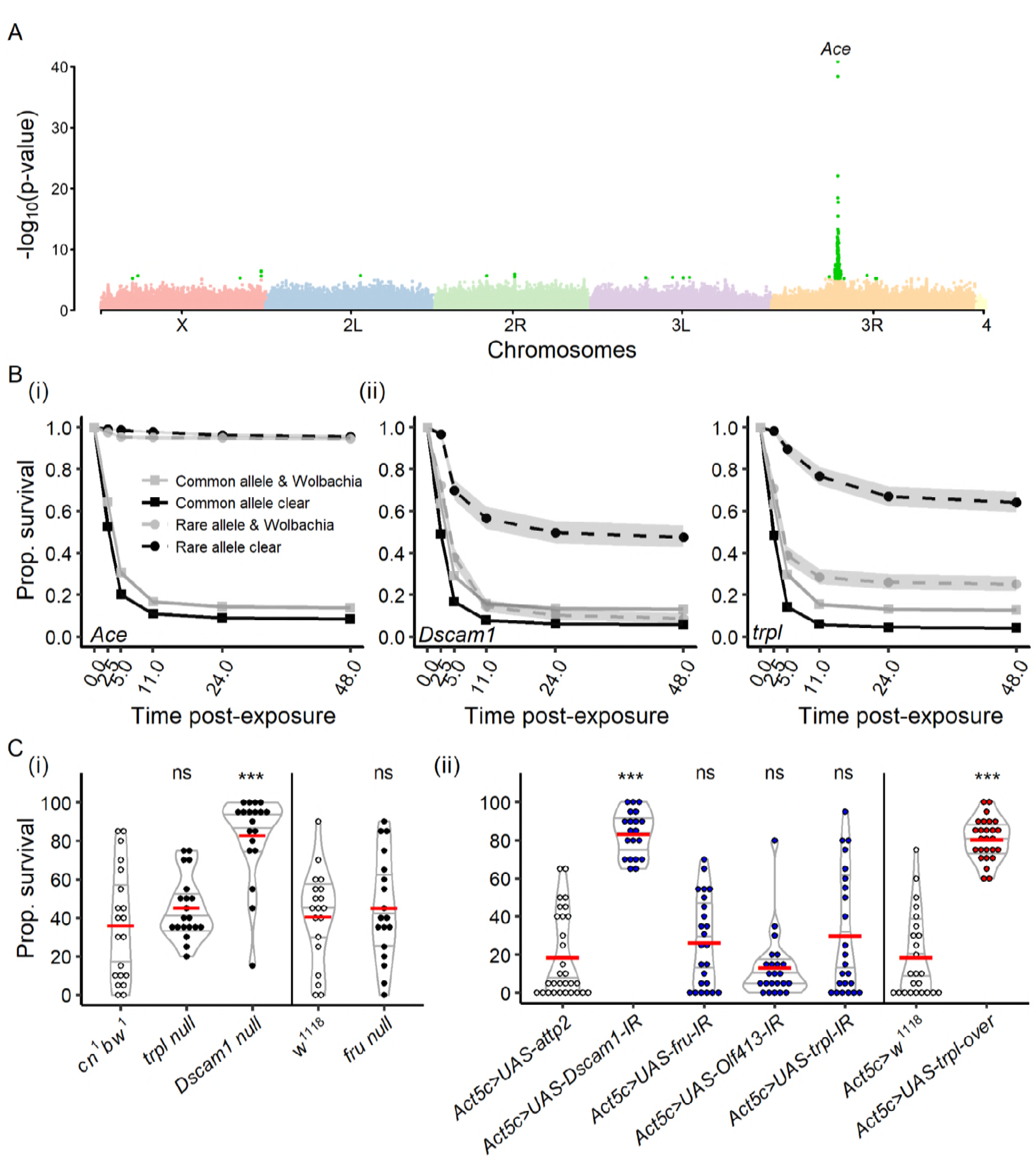
**A**-Manhattan plot describing the results of the main GWAS on parathion resistance (including 194 DGRP lines). Light green dots represent the SNPs with a p-value below a 10^-5^ threshold. Loci in the *Ace* gene were the main loci responsible for the variation in resistance to parathion exposure. **B**-Survival curves (in hours) of lines variants for the validated candidate genes for resistance to parathion. *i*) Variation in *Ace* (mutation F368Y) in position 3R:13,243,332 affects the resistance to parathion. *ii*) Variation in *Dscam1* affects the resistance to parathion, but only in lines that do not carry *Wolbachia* (Survival analysis with lognormal distribution: interaction SNP and *Wolbachia*: deviance= 455.39, p < 0.0001). *iii*) Variation in *trpl* affects the resistance to parathion, but only in lines that do not carry *Wolbachia* (Survival analysis with lognormal distribution: interaction SNP and *Wolbachia*: deviance= 735.69, p < 0.0001). **C**-Validation of the candidate genes of our GWAS. White dots represent the wildtype genotypes, black dots the loss-of-function mutants, blue dots the downregulation and red dots the upregulation of the genes. Non-significant effects are indicated by “ns”, p-values below 0.001 are indicated by ***. Details of the statistics are summarized in Table S6 and S7.

The dominant role of *Ace* SNPs in causing resistance to parathion presented the potential for this strong signal to mask other genes involved in resistance (e.g. those with a lower effect).

To identify these secondary loci associated with parathion resistance, we next performed a nested GWAS. For that purpose, we ran a new GWAS using only a subset of lines (n= 124) that did not carry the resistance allele for the most significant SNP (i.e. mutation F368Y in the *Ace* gene). This association was tested over 1,212,116 remaining SNPs/indels. Amongst those, we identified a list of candidates with the same criterion as above (grey p-values in Figure S4, Table S3). From this list, we selected four candidate genes based on the annotated function of the protein and the availability in stock centers of genetic tools to perform functional validation: *trpl* (Figure 2B*ii*) that encodes a non-selective cation channel, *olf413* that encodes a dopamine beta hydrolase, *fru* that encodes a key determinant of sex specific expression, and *Dscam1* (Figure 2B*ii*) that encodes a transmembrane receptor involved in neuron wiring. The mutations in the genes coding for *Dscam1* and *trpl* were only associated to an increase in resistance with lines not infected by *Wolbachia* [Figure 2B*ii* (*Dscam1*), Survival with lognormal distribution: interaction SNP and *Wolbachia*: deviance= 455.39, p< 0.0001; Figure 2B*ii* (*trpl*), Survival with lognormal distribution: interaction SNP and *Wolbachia*: deviance= 735.69, p< 0.0001]. This result suggests strongly that *Wolbachia* could have a direct role in the resistance to insecticides, but this effect depends on host genotype. Alternatively, it is possible that *Wolbachia*’s presence alters the activity of other unidentified genes involved in resistance. We next analyzed the impact of loss of function (null) alleles or RNAi knockdown of these candidate genes on the susceptibility to parathion. RNAi-mediated knock-down of *olf413* or *fru* expression did not result in any changes in survivorship, suggesting they are not involved in resistance to parathion (Figure 2C). However, both downregulation of *Dscam1* by RNAi and a null mutation of *Dscam1* confirmed its role in resistance to parathion (Figure 2C). Knock-down of *trpl* did not affect susceptibility to parathion, but upregulation of *trpl* strongly increased resistance to parathion (Figure 2C).

Overall, our results strongly suggest that *Ace*, *Dscam1* and *trpl* are important for resistance to parathion and are involved in the phenotypic variation between strains. A possible mechanism by which these genes could contribute to resistance would be due to changes in their constitutive expression. To test this, we took advantage of a previous study that measured the expression of transcripts genome-wide in the DGRP lines (Huang *et al*. 2015). There was no correlation between constitutive expression of *Ace*, *Dscam1* and *trpl* in the conditions of their study and our survival experiments (Figure S6A-C). Altogether, our data demonstrate that the genetic basis for the variation in resistance to parathion is multigenic, with a major effect due to non-synonymous mutations in *Ace* and secondary roles due to mutations in *Dscam1* and *trpl* that can be buffered by the presence of *Wolbachia*.

### Genetic basis of the variation in resistance to deltamethrin

Using the same strategy outlined above, we analyzed the association of 2,171,433 SNPs/indels with deltamethrin survival. In total, 6 loci were strongly significantly associated (*i.e.* -log_10_(p-value) > 8) to resistance to deltamethrin at the 48h time point but other, less strongly associated, SNPs/indels could be considered as potential candidates (192 had -log_10_(p-value) > 5 and 1066 had -log_10_(p-value) > 4) (Figure 3A, Figure S7, Table S4). Among the most significant, two non-synonymous mutations strongly associated with resistance to deltamethrin were mapped to *Cyp6a23* (Figure 3B, 2R:14,876,125; Figures S8A, 2R:14,876,857). The peak of association was detected in *Cyp6a23.* However, there are five other *Cyp*s at this locus (Figure 3C) and few SNPs in non-coding or intergenic regions were significantly associated with resistance within this locus (Figure 3A inlet). Thus, we wanted to test the possibility that other *Cyps* in the locus might also be involved in resistance to deltamethrin (no missense SNPs/indels in any of the other *Cyps* of the locus were significantly associated to resistance, but the information of the SNPs/indels is incomplete). We therefore decided to test all six *Cyps* (*Cyp6a23*, *Cyp6a9*, *Cyp6a19*, *Cyp6a20*, *Cyp6a17* and *Cyp6a22)* using all the available RNAi lines against these *Cyp* genes and using the one null line (*Cyp6a17*) available. Knocking down *Cyp6a23* and *Cyp6a17* increased susceptibility of flies to deltamethrin (Figure 3D*i*). In contrast, but not so surprisingly (based on Figure 3A inlet), knocking down the other *Cyp*s did not change the survival to deltamethrin in comparison to their genetic control (Figure 3D*i*; Figure 3D*ii*). We further confirmed the role of *Cyp6a17* in resistance to deltamethrin by using a null mutant (Figure 3D*iii*). These results imply that only two *Cyp* genes in that locus are involved in resistance to deltamethrin: *Cyp6a23* (major effect) and *Cyp6a17* (secondary effect), although we do not know whether there are any mutations in *Cyp6a17* that could provide resistance. Remarkably, these two neighboring genes are paralogous (Figure 3C) (*i.e.* two genes descend from a common ancestral DNA sequence and derive within one species) (Good *et al*. 2014) and reminds us of *Ace-1* and *Ace-2*, two homologous genes involved in insecticide resistance in mosquito species (Weill *et al*. 2002). Cyp-mediated resistance can occur through changes in gene expression (Liu and Scott 1998) or structural changes (Amichot *et al*. 2004). Therefore, we next asked whether DGRP flies expressed different levels of *Cyp6a23* and *Cyp6a17*, and whether these expression levels correlated with resistance. The constitutive expression of *Cyp6a23* estimated in (Huang *et al*. 2015) did not correlate with a higher resistance to deltamethrin (Figure S6D). However, there was a strong positive correlation with the constitutive expression of *Cyp6a17*, consistent with our results (Figure S6E).

**Figure 3.**
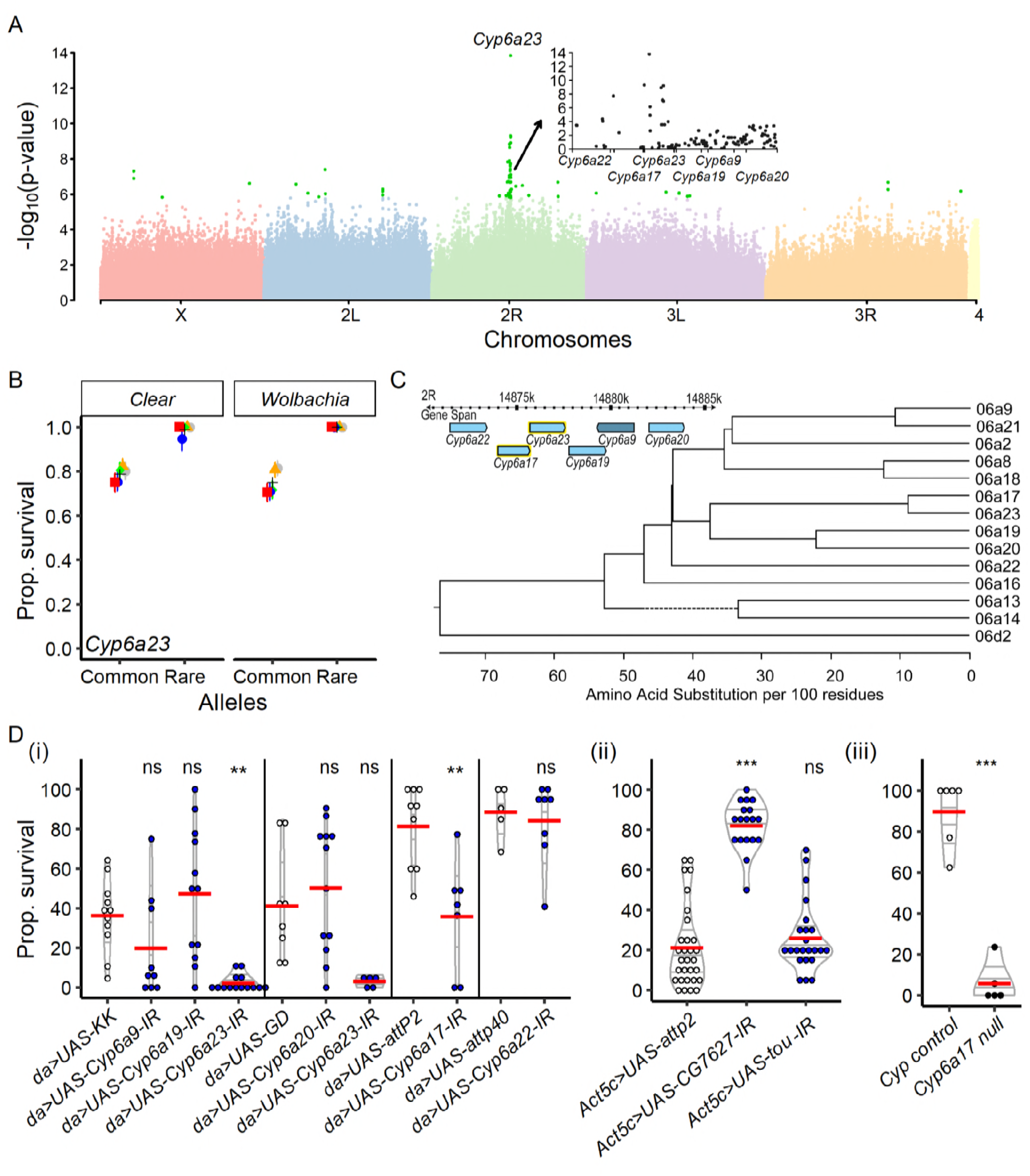
**A**-Manhattan plot describing the results of the main GWAS on deltamethrin resistance (including 195 DGRP lines). Light green dots represent the SNPs with a p-value below a 10^-5^ threshold. The locis mainly responsible for the variation in resistance to deltamethrin exposure were located in the *Cyp6a23* gene or its direct proximity, within the *Cyp6a* cluster. Inlet graph represents a magnification of the results and suggests that **Cyp6a*23* and *Cyp6a*17 were the most likely candidates. **B**-Mean survival of lines variants for the validated candidate genes **Cyp6a*23* for resistance to deltamethrin. Colors represent five replicated experiments. **C**-*Cyp6a*23 is part of a cluster of genes belonging to the cytochrome P450 family. The phylogeny represents the already suggested hypothesis that **Cyp6a*23* and *Cyp6a17* are two neighboring paralogous genes issued from a recent duplication. **D**-Validation of the candidate genes of our GWAS. White dots represent the wildtype genotypes, black dots the loss-of-function mutants and blue dots the downregulation of the genes. Non-significant effects are indicated by “ns”, p-values below 0.01 are indicated by ** and p-values below 0.001 are indicated by ***. Details of the statistics are summarized in Table S6 and S7.

**Figure 4.**
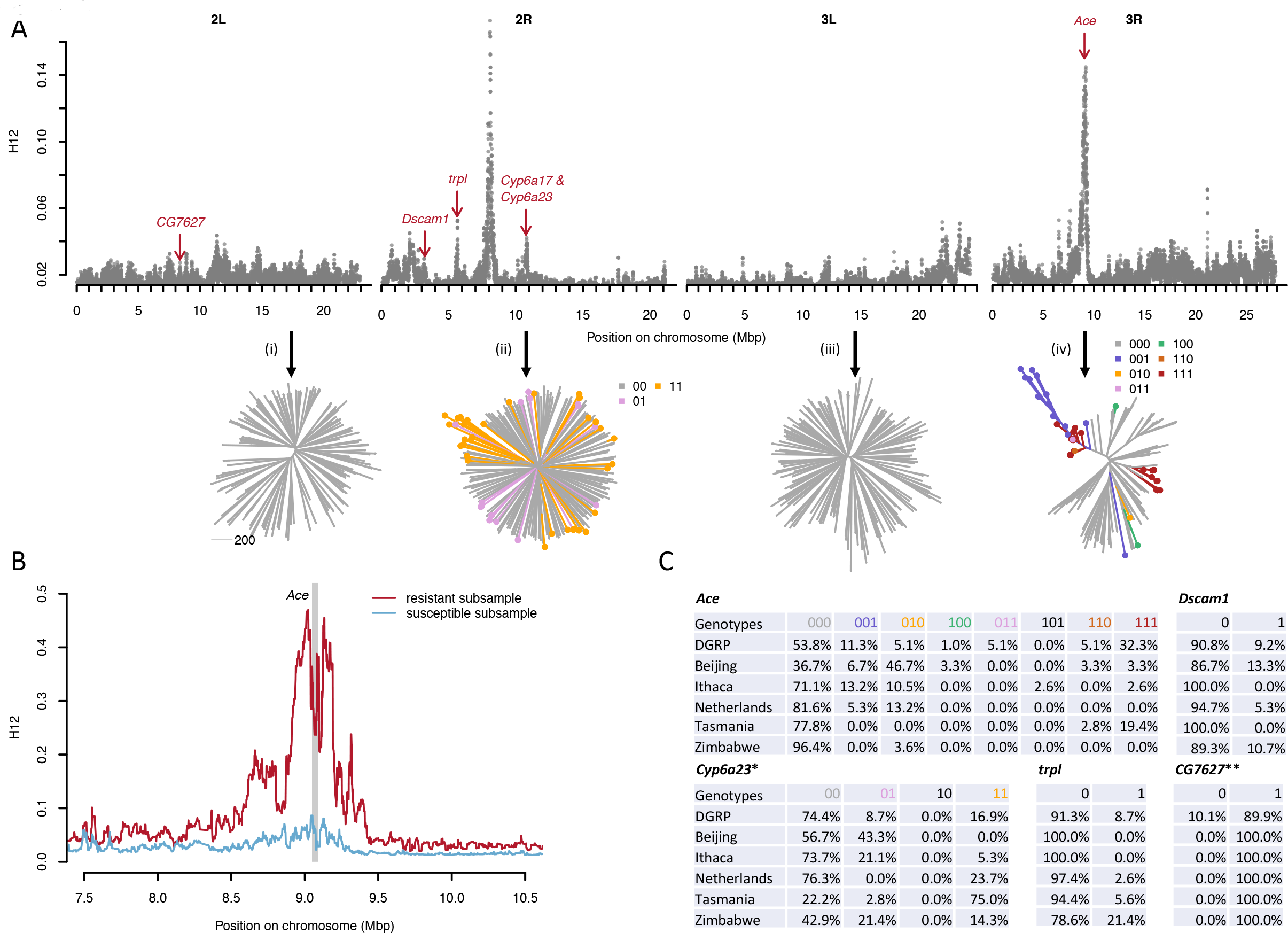
**A**-Genome-wide H12 scan for all autosomal SNPs in the DGRP data, using window sizes of 800 segregating sites centered around each focal SNP. Red arrows indicate the positions of our candidate loci. The lower panel shows neighbor-joining tress for selected genomic windows of length 200 kbp from each autosomal arm: (i) a random window on 2R, (ii) window centered on the *Cyp6a23* locus, (iii) a random window on 3L, and (iv) a window centered on the *Ace* locus. The coloring of the leaf nodes in (ii) and (iv) specifies the particular combination of resistance mutations each haplotype carries at the respective locus (e.g. 011 indicating presence of the second and third resistance mutation at *Ace*, while 000 indicates a haplotype with none of the three resistance mutations). **B**-H12 scan around the *Ace* locus after splitting the DGRP data into two subsets of genomes that either carry at least one of the three resistance mutations (resistant haplotypes) or do not carry any such mutation (susceptible haplotypes). The latter group was down-sampled so that both subsamples comprised the same number of genomes (n = 90). **C**-Frequencies of resistance mutations in the DGRP data and the five-continent reference panel of the global diversity lines (GDL) (Grenier *et al*. 2015). *In Zimbabwe, at the first **Cyp6a23** resistance locus an alternative allele is present in ∼21.4% of the GDL strains that is not found in the DGRP, and for which we therefore do not know whether it is a resistant or susceptible allele. **At the *CG7627* locus, the resistant allele is the reference allele and the susceptible allele is an insertion of a single base pair. We did not observe this insertion in any of the GDL lines (although it could be possible that this indel exists in the panel but was not called in the data).

To identify secondary loci associated with deltamethrin resistance, we performed a nested GWAS using only a subset of lines (n= 147) that did not carry the resistance allele for the most significant SNP (*i.e.* in position of 2R:14,876,125 of *Cyp6a23*). The association was tested over 1,872,071 SNPs and we identified 11 SNPs/indels significantly associated (-log_10_(p-value) > 8), 142 with a -log_10_(p-value) > 5 and 766 with a -log_10_(p-value) > 4 with resistance against deltamethrin (Table S5). Among the significant SNPs/indels, an isolated indel with a high p-value (-log_10_(p-value) = 6.44, Figure S8B) was close and upstream from the gene *CG7627,* which appears to have ATPase activity and be involved in transmembrane movement of substances. Flies in which we downregulated the expression of *CG7627* by RNAi had a lower probability to die from the exposure to deltamethrin when compared to their control (Figure 3D*ii*), although the constitutive expression of this gene did not correlate with resistance (Figure S6F). We also tested the role of *toutatis* (*tou*) which interestingly was associated with resistance to deltamethrin in both the GWAS and nested GWAS (Figure S7) and is supposedly involved in nervous system development (Vanolst 2005). However, the knock-down of this gene by RNAi did not confirm a role of this gene in resistance (Figure 3D*ii*). This might not be surprising as the change associated to resistance was a synonymous mutation in an intronic region of the gene (Table S5).

Overall, we find that deltamethrin resistance is primarily due to non-synonymous mutations in *Cyp6a23* and increased expression of *Cyp6a17*. RNAi of *Cyp6a23* suggests this gene is capable of detoxifying deltamethrin, yet no correlation of *Cyp6a23* constitutive expression (estimated in Huang *et al*. 2015) and deltamethrin survival was found. RNAi and null strains suggest that *Cyp6a17* is capable of detoxifying deltamethrin and the constitutive expression estimated in (Huang *et al*. 2015) of *Cyp6a17* correlates with deltamethrin survival, yet the GWAS signal is not centered over *Cyp6a17*. We validated *CG7627* as having a secondary effect on survivorship.

### Loci associated with resistance to insecticides show signatures of positive selection

We found that a small number of individual loci explain most of the variation in resistance across the DGRP lines for both parathion and deltamethrin, suggesting that these loci could have undergone recent positive selection. To test this hypothesis, we performed a genome-wide scan of the DGRP lines using the H12 statistic (Garud *et al*. 2015). This statistic estimates levels of haplotype homozygosity and has previously been shown to provide good power in detecting both hard and soft selective sweeps (Garud *et al*. 2015; Miles *et al*. 2016). A previous H12 scan of the DGRP has already detected a strong sweep signal at the *Ace* locus, as well as two other loci known to be associated with insecticide resistance (*ChKov1* and *Cyp6g1*) (Garud *et al*. 2015; Schmidt *et al*. 2017). Our genome-wide scan presented in Figure 4A confirms these signals and also reveals clear sweep signatures at all of the other key resistance loci identified in our GWAS analysis (*CG7627*, *Dscam1*, *trpl*, and *Cyp6a23*/*Cyp6a17*). Many of these signals rank among the most pronounced sweep signals detected genome-wide, suggesting that the evolution of pesticide resistance constitutes one of the strongest adaptive response experienced by *D. melanogaster* in its recent evolutionary history.

### Haplotypes at *Ace* are consistent with a soft selective sweep driven by resistance alleles

To demonstrate that the signals of positive selection we observed in the genome-wide H12 scan were indeed driven by the specific resistance mutations, rather than some other alleles, we studied patterns of haplotype diversity at several resistance loci using neighbor-joining trees (Figure 4A). The haplotype tree around *Ace*, which constituted the strongest signal in the H12 scan, showed clear signatures that the sweep patterns observed at this locus were indeed driven by the resistance mutations, as indicated by the presence of several independent clusters of resistance mutation-carrying haplotypes with short genetic distances within clusters. Susceptible haplotypes, by contrast, showed patterns similar to the genomic background. In particular, we observed two distinct clusters of haplotypes carrying resistance mutations at all three sites (111). One of these clusters is located close to a cluster of haplotypes carrying only the third resistance mutation (001), suggesting a short evolutionary distance between these haplotypes. All haplotypes we observed in the DGRP that carried resistance mutations at two of the three sites (011 & 110) also fell in this group. This is consistent with a scenario in which these two-mutation haplotypes represent transition haplotypes to three-mutation haplotypes, or back-mutations. We observed several low-frequency haplotypes with only one resistance mutation (100, 010, and 001) that did not appear to cluster with any of the other resistance haplotypes, suggesting that these haplotypes arose independently from wildtype alleles, as has been proposed previously (Karasov *et al*. 2010).

To provide further evidence that the sweep signal at *Ace* is indeed driven by the resistance mutations, we split the DGRP lines into two subsamples, the first comprising the genomes that carry at least one of the three resistance mutations, and the second comprising those that do not carry any such mutation. We then estimated H12 independently in each subsample (after down- sampling the second sample to the same size as the first). Figure 4B shows that the H12 peak is only observed in the subsample with resistance mutations, whereas there is almost no such signal among the susceptible genomes. This again confirms that it is indeed the resistance mutations (or some very tightly linked mutations) that primarily drive the peak in the H12 signal around *Ace*.

At the *Cyp6a23*/*Cyp6a17* loci we also detected sweep signatures in our H12 scan, although these signals were much weaker than at the *Ace* locus. One possible explanation for this is that the *Cyp6a* locus has undergone a very soft sweep from standing variation, which is consistent with the fact that the haplotype tree at this locus does not show any noticeable clustering of resistance alleles (Figure 4A). In addition, the resistance mutations are at very low frequency at the *Cyp6a23*/*Cyp6a17* locus in the DGRP data, limiting the extent of possible sweep signatures.

### Global distribution of resistance allele frequencies

To study the global prevalence of the different resistance mutations identified in our GWAS we estimated their frequencies in the DGRP, as well as a panel of Global Diversity Lines (GDL) comprising fly strains from five different continents (Grenier *et al*. 2015). Figure 4C shows the frequencies of resistant (1) and susceptible (0) alleles — and combinations thereof at individual loci — for *Ace*, *Cyp6a23*, *Dscam1*, *trpl*, and *CG7627*, revealing substantial frequency variation between populations. For example, haplotypes with neither of the two resistance mutations at the *Cyp6a23* locus (00) constitute only ∼22% of the strains from Tasmania, but ∼74% of the DGRP strains. By contrast, fully resistant strains (11) constitute ∼75% of the strains from Tasmania, yet only ∼17% in the DGRP. These patterns could suggest that more intense pyrethroid selection has occurred in Tasmania compared to the rest of the world. Allele frequency differences are even more pronounced at *Ace*. Here, haplotypes with none of the three resistance mutations (000) comprise ∼96% of the strains from Zimbabwe, but only ∼37% of strains from Beijing, suggesting that the least intense organophosphate selection has occurred in the Zimbabwe population. Among the resistant haplotypes at *Ace*, there is also surprising variation in terms of the frequencies of individual resistance allele combinations. For instance, the most common combination of resistance alleles in the DGRP is 111 at ∼32%. Most of the other possible configurations with one or two resistance mutations also occur, yet at much lower frequencies. In the Beijing sample, however, the most frequency resistant configuration is 010 at ∼47%, with the three-mutation configuration (111) present in only ∼3% of strains. This extensive diversity in resistant haplotypes is consistent with a non-mutation-limited scenario in which individual resistance mutations can evolve rapidly and repeatedly at individual loci, such that even complex, multi-step adaptations can arise quickly with intermediate configurations not necessarily reaching high population frequency (Messer and Petrov 2013). This is also consistent with the possibility that different insecticides (carbamates and/or structurally different OPs) were used in different regions and that they are selecting for different mutations (Oppenoorth 1985).

## Discussion

The evolutionary outcome from insecticide selection has proven to be extraordinarily difficult to predict and our results confirm this. We find that the results with deltamethrin were very unexpected, as no changes in the target site gene were found. This is in stark contrast to both how pyrethroid resistance has evolved in most insects, and to parathion where most of the resistance was conferred by *Ace* mutations. Furthermore, the genes identified and validated as having a secondary role in resistance to parathion or deltamethrin would not have been the ones that were expected based on previous resistance work. However, there were some consistencies between the parathion and deltamethrin results. The most notable part is that most of the resistance in both cases was primarily due to mutations at a single locus. The debate over whether insecticide resistance is most commonly monogenic or polygenic will not easily be resolved, as there are clear examples that both occur. Our data suggest that resistance to parathion and deltamethrin in the DGRP lines are polygenic, but that a single locus confers most of the resistance.

Much of the work on insecticide resistance has focused on changes in target site or detoxification genes, in part for historical reasons. However, identification of other genes that can be involved in resistance has been very challenging. GWAS studies like what we did have the potential to identify toxicologically relevant genes that would otherwise be very difficult to identify. For example, our studies implicate *Dscam1* and *trpl* in parathion resistance and *CG7627* in resistance to deltamethrin. Based on what is known about these genes it is difficult to provide a physiological or toxicological explanation for their role. However, these are exciting genes for further investigations that could greatly improve our understanding of the poisoning process in insects. The former, the *Down syndrome cell adhesion molecule 1* (*Dscam1*) is known for its involvement in self-avoidance mechanisms that are key during neurogenesis. It is not entirely surprising that it plays a role in the resistance against an insecticide that disrupts the nervous system. The later, *CG7627*, is known to be involved in membrane transport. We do not know much about this gene, but other proteins that are capable of transporting xenobiotics can alter the toxicity of insecticides (Sun *et al*. 2017). Most genetic variance for resistance relies on genes with a major effect, however, other genes clearly play a significant role.

Surprisingly, the genetics of resistance can be altered by the presence of *Wolbachia*. Beyond the fact that GWAS generally ignores the epistatic effect among genes, our study reveals clearly that the effect of resistant alleles can depend on *Wolbachia* infection. *Wolbachia* density can correlate positively with the presence of insecticide-resistant genes in mosquitoes (Berticat *et al*. 2002), however, it seems that the pleiotropic effect of *Wolbachia* on resistance alleles can have a major influence on the efficiency of the resistance, as it is the case for *Dscam1* and *trpl*. This implies that *Wolbachia* could be a buffer to the effect of resistance alleles and prevent them from fixation.

Fruit production relies heavily on the use of insecticides. As such, *D. melanogaster* is expected to be under a strong selection pressure to develop resistance. Our results confirm this happening in the field, particularly for OPs and pyrethroids which were used in the decades preceding the collection of the DGRP lines. We selected parathion and deltamethrin as our prototypical OP and pyrethroid, respectively. However, what we observed in the DGRP lines is not necessarily the result of exclusive selection with parathion or deltamethrin, but rather the combined results of all OPs (and carbamates) and pyrethroids. This is important simply to prevent over-interpretation of our results. For example, the mutations in *Ace* that resulted in parathion resistance in the DGRP lines are likely the result of cumulative selection with multiple OPs (and carbamates), not necessarily the result of selection only with parathion. Conversely, *Cyp6a23* is not involved in resistance to DDT, nitenpyram, dicyclanil nor diazinon (Daborn *et al*. 2007), but the selection on this gene could be due to pyrethroids other than deltamethrin.

While it is remarkable that the GWAS analysis for both insecticides identified a single locus, it is curious that in one case variation in toxicity was linked to mutations in the target site gene (*Ace* for parathion), but not for the other (*Vss*c for deltamethrin). This is not limited to the DGRP lines as evaluation of the Global Diversity Lines also showed that mutation in *Vssc* was not present. This makes *D. melanogaster* quite unusual as *Vssc* mutations are very common in pest species and have been found in at least one strain from virtually every pyrethroid/DDT resistant species examined (Dong *et al*. 2014). One possibility would be if there was a codon usage in *D. melanogaster*, such that the resistance mutation could not occur with a single nucleotide change. This has been proposed as a reason why organophosphate and carbamate insecticides had not selected for the G119S mutation in *Ace* in *Aedes aegypti* (Weill *et al*. 2004). The most common *Vssc* mutation is L1014/F/H/S/C/W (house fly numbering system) (Scott *et al*. 2013). The codon used by *D. melanogaster* at this position is CTT (same as house fly). Thus, a single nucleotide change could produce known resistance mutations at this position. Similarly, the T929I mutation can also confer pyrethroid resistance (Dong *et al*. 2014) and the codon at this position in *D. melanogaster* could accommodate this change with a single nucleotide mutation (from ACA to ATA). However ethyl methanesulfonate (EMS) mutagenesis led to the recovery of *para* (the *D. melanogaster Vssc*) mutants that were up to 22-fold resistant to DDT, and up to 10-fold resistant to deltamethrin (Pittendrigh *et al*. 1997) and recently the I265N *para* mutation was found to confer 6.3-fold resistance to deltamethrin (Rinkevich *et al*. 2015). In contrast, permethrin selection of wild caught *D. melanogaster* failed to generate a resistant strain (R. Roush, personal communication), although cyclodiene selection of the same populations was highly successful (ffrench-Constant *et al*. 1990). Thus, under laboratory conditions *para* mutations can be made that result in insensitivity to pyrethoids (and DDT), but such mutations do not appear to underlie resistance in field populations of *D. melanogaster* (based on the DGRP and GDL lines and laboratory selections of field populations). It is difficult to reconcile why selection favored changes in a target site for OPs and yet favored changes in a detoxification gene for pyrethroids.

Our results provide an interesting comparison to the three other papers that have evaluated the DGRP lines to look for loci associated with resistance to DDT, azinphos-methyl and imidacloprid (Battlay *et al*. 2016; Schmidt *et al*. 2017; Denecke *et al*. 2017). Most striking is that different genes are responsible for azimphos-methyl and parathion, even though both are OPs. The major gene associated with azinphos-methyl resistance was *Cyp6g1* with a secondary effect seen for *CHKov1* (Battlay *et al*. 2016). In contrast, the major gene associated with parathion resistance was *Ace* with secondary effects seen for *Dscam1* and *trpl*. Although mutations in *Ace* are a common mechanism of resistance to OPs (and carbamates), it has long been recognized that mutations in *Ace* that give insensitivity to one insecticide may provide little or no resistance to other OPs (or carbamates) (Oppenoorth 1985). However, the *Ace* mutations present in the DGRP lines render the protein less sensitive to inhibition by azinphos-methyl oxon, the bioactivated form of azinphos-methyl (Menozzi *et al*. 2004). One possibility why *Ace* was not detected as a locus for resistance to azinphos-methyl would be if *Cyp6g1* was highly efficient at detoxification of this insecticide, such that the bioactivated form was not produced in lines that had this resistance allele. However, the *Ace* and *Cyp6g1* mutations would be expected to segregate, giving a signal for both mutations and making it unclear why this locus was not detected for azinphos-methyl resistance (Battlay *et al*. 2016).

DDT was widely used from 1946 until resistance problems became wide spread (about 1960) and other more effective insecticides were introduced. DDT was banned by EPA in 1972. Organophosphates were introduced in the mid-1940s and became the most widely used class of insecticides from about 1955 – 1987. Pyrethroids were introduced about 1980 and rapidly rose to become the most widely used class of insecticides from about 1989-2000. Neonicotinoids (specifically imidacloprid) was registered for use in fruit about 1994 and have been the most widely used class of insecticides since about 2000. The DGRP lines were collected in 2003 (Mackay *et al*. 2012). Thus, use of the DGRP lines to evaluate DDT resistance would be searching for signs of selection that would have ceased nearly 50 years ago. In the case of OPs and pyrethroids, the selection has been ongoing for over 50 and 30 years, respectively. In the case of neonicotinoids, the selection would have been for only about a decade. Based on this, we might expect that we would detect the strongest to weakest signals for parathion, followed by deltamethrin and then imidacloprid and/or DDT. Exceptions to this might occur if there was cross-resistance between one of these insecticides and what was used in the field. Given the different loci that were detected for parathion, deltamethrin and imidacloprid, suggests this is unlikely and indicates the detected loci were the result of OP or carbamate, pyrethroid and neonicotinoid insecticides, respectively. However, *Cyp6g1* was detected for DDT, azinphos-methyl and imidacloprid resistance. Thus, the GWAS analysis for DDT may not represent what evolved in the population due to DDT use, but rather what evolved in the population over the last 40 years that conferred cross-resistance to DDT.

Altogether our study confirms that insecticides apply a strong selection pressure even on insects, like *D. melanogaster*, that are not the targeted pest and highlight that pesticide management should take into account the effect on the whole insect community. Furthermore, the fact that resistance can be buffered by the presence of the common endosymbiont *Wolbachia* and can evolve through changes in target site or in detoxification enzyme depending on the insecticides and on the insect species make evolution of resistance in those communities fairly unpredictable. However, resistance alleles were present in populations sampled throughout the world showing that even if unpredictable, evolution of resistance to insecticide is repeatable.

## Acknowledgements

We thank Jean-Baptiste Ferdy and Fabrice Roux for thoughtful discussions and Pierre Solbes for support with the cluster. This project was supported by start-up funds (to NB). HS was supported by a fellowship from the China Scholarship Council (Grant No. 201406850044). DD was supported by the French Laboratory of Excellence project “TULIP” (ANR-10-LABX-41; ANR- 11-IDEX-0002-02) and by the People Programme (Marie Curie Actions) of the European Union’s Seventh Framework Programme (FP7/2007-2013) under REA grant agreement n. PCOFUND-GA-2013-609102, through the PRESTIGE programme coordinated by Campus France. HDK and IVC were supported by Presidential Life Science Fellowships from Cornell University. Computations were performed on the EDB-Calc Cluster which uses a software developed by the Rocks(r) Cluster Group (San Diego Supercomputer Center, University of California, San Diego and its contributors), hosted by the laboratory “Evolution et Diversité Biologique” (EDB).

## Supplementary tables

**Table S1** List of *D. melanogaster* genotypes. Stock number refers to the Bloomington or VDRC stock center numbers.

**Table S2** Results of GWAS of parathion resistance.

**Table S3** Results of nested GWAS of parathion resistance.

**Table S4** Results of GWAS of deltamethrin resistance.

**Table S5** Results of nested GWAS of deltamethrin resistance.

**Table S6** Details of the validation (see Figure 2C and 3D). Results from general linear hypothesis test (glht) with Tukey post Hoc pairwise comparisons, to ascertain differences between pairs of treatments (package *multcomp* in R) after a generalized linear model with a quasibinomial distribution of the residuals.

**Table S7** Details of the validation (see Figure 2C and 3D). Results from generalized linear model with a quasibinomial distribution of the residuals.

**Table S8** Raw phenotypic data for resistance to Parathion.

**Table S9** Raw phenotypic data for resistance to Deltamethrin.

## Supplementary figure legends

**Figure S1** Difference in survival to insecticide exposure between the DGRP lines carrying *Wolbachia* and those that do not carry the endosymbiont. Lines carrying *Wolbachia* did not survive better than those without *Wolbachia* (A: Survival to parathion over time; B: Survival to deltamethrin at 48 h). Non-significant effects are indicated by “ns”.

**Figure S2** Correlation between the resistance to insecticide (i.e. proportion surviving after 48 h of parathion or deltamethrin exposure) and other abiotic stresses: Paraquat (**A** and **B**), Starvation (**C** and **D**) and alcohol (alcohol sensitivity is measured by measuring elution time) (**E** and **F**). Measurements of resistance to other stresses were performed in other studies (see details in methods). Analysis of correlation was done with Spearman correlation test. A blue line represents the significant correlation between the two traits.

**Figure S3** Genetic correlation between 10 lines amongst the most sensitive (red) and 10 lines the amongst most resistant (green) to **A**) parathion exposure and **B**) deltamethrin exposure. The grey gradient represents the strength of the genetic correlation with black being “genetically identical”.

**Figure S4** Manhattan plots with the package *Chromplot* in R showing precisely the peak of p-values along the genome for the complete GWAS (shown to the left of the chromosome) and the nested GWAS (shown to the right of the chromosome) for resistance to parathion. Names of genes are manually selected candidates. The full datasets can be found in tables S2 and S3.

**Figure S5** Mean survival upon parathion exposure of lines variants at the *Ace* loci. **A**- Variation in *Ace* (mutation G303A) in position 3R:13,243,686 affects the resistance to parathion. **B**- Variation in *Ace* (mutation I199V) in position 3R:13,243,999 affects the resistance to parathion.

**Figure S6** Correlation between the resistance to insecticide (i.e. proportion of survival 48 h upon parathion or deltamethrin exposure) and the constitutive expression of validated genes. Experimental measurements of gene expression were measured in other studies (see details in methods). Analysis of correlation was done with Spearman correlation test. A blue line represents represent the significant correlation between the two traits.

**Figure S7** Manhattan plots with the package *Chromplot* in R showing precisely the peak of p-values along the genome for the complete GWAS (shown to the left of the chromosome) and the nested GWAS (shown to the right of the chromosome) for resistance to deltamethrin. Names of genes are manually selected candidates. The full datasets can be found in tables S4 and S5.

**Figure S8** Mean survival upon exposure to deltamethrin of lines variants for the SNP in position 2R:14,876,857 belonging to the validated candidate gene *Cyp6a23* (**A**) and for the SNP belonging to the validated candidate gene *CG7627* (**B**). Colors represent five replicated experiments.

